# Training-induced alterations in the modulation of human motoneuron discharge patterns with contraction force

**DOI:** 10.1101/2025.06.02.657380

**Authors:** Jakob Škarabot, Haydn W Thomason, Benjamin M Nazaroff, Christopher D Connelly, Tamara Valenčič, Michael L Ho, Kapil Tyagi, James A Beauchamp, Gregory EP Pearcey

## Abstract

Motoneurons adapt to both resistance and endurance training in reduced animal preparations, with adaptations seemingly more apparent in higher threshold neurons, but similar evidence in humans is lacking. Here, we compared the identified motor unit (MU) discharge patterns from decomposed electromyography signals acquired during triangular dorsiflexion contractions up to 70% of maximal voluntary force (MVF) between resistance-trained, endurance-trained, and untrained individuals (n=23 in each group). We then estimated intrinsic motoneuron properties and garnered insight about the proportion of excitatory, inhibitory, and neuromodulatory inputs contributing to motor commands across contraction intensities in each group. Participants also performed a task where a triangular contraction was superimposed onto a sustained one designed to challenge inhibitory control of dendritic persistent inward currents (PICs). Both trained groups demonstrated greater MU discharge rates with greater ascending discharge rate modulation during higher contraction forces (≥50% MVF), which were accompanied by more linear MU discharge patterns and greater post-acceleration attenuation slopes of the ascending discharge rates. No differences in discharge rate hysteresis or the discharge rate characteristics during the sombrero tasks between groups, suggesting no differences in neuromodulatory input. Conversely, resistance-compared to endurance-trained individuals exhibited greater acceleration slopes during lower contractions forces (≤50% MVF), indicating the possibility of enhanced initial activation of PICs. Collectively, the greater and more linear MU discharge patterns in the trained groups either suggests a more reciprocal (i.e., push-pull) excitation-inhibition coupling during higher contraction forces or enhanced excitatory synaptic input to the motor pool, which might underpin greater force production of trained individuals.

## INTRODUCTION

Motoneurons are highly adaptable to a variety of stressors, including exercise (for reviews, see Gardiner et al., 2006; Macdonell & Gardiner, 2018; Pearcey et al., 2021; Škarabot et al., 2020). From an electrophysiological perspective, endurance exercise in rodents has been shown to hyperpolarise motoneuron resting membrane potential and voltage threshold, and decrease the spike rise time, the threshold for rhythmic discharge, the minimal steady-state discharge frequency, and the slope of the frequency-current relationship (Beaumont & Gardiner, 2002, 2003). Similarly, motoneuron adaptations have been shown in rodent models following resistance training, including an increase in the time course of afterhyperpolarisation and input resistance, and a decrease in minimal steady-state discharge frequency and rheobase current (Krutki et al., 2017). These changes in intrinsic motoneuron discharge properties with exercise have been attributed to altered ion conductance, particularly those of sodium channels (Gardiner et al., 2006). However, recent evidence suggests prominent exercise-induced dendritic plasticity leading to alterations within the voltage-gated sodium and calcium channels facilitating dendritic persistent inward currents (PICs) to interneurons (Chen & Dai, 2022; Ge & Dai, 2020). PICs are also critical for modulating intrinsic excitability of motoneurons and are facilitated by diffuse metabotropic/neuromodulatory inputs via long descending axons of the raphe nuclei and locus coeruleus (Bennett et al., 1998; Goaillard & Marder, 2021; Hounsgaard et al., 1988; Schwindt & Crill, 1980). Due to the diffuse nature of metabotropic inputs, the gain control of motoneuron output is mediated by the potency and pattern of inhibitory inputs to supress PICs (Hyngstrom et al., 2007; Johnson et al., 2012; Pearcey et al., 2022; Revill & Fuglevand, 2017). During a linearly increasing excitatory synaptic input, PICs introduce non-linearities in motoneuron discharge rate by amplifying and prolonging the excitatory input (Lee & Heckman, 1998, 2000), allowing the estimation of intrinsic motoneuron properties and the proportion of excitatory, inhibitory, and neuromodulatory motor commands from human motor unit (MU) discharge patterns (Beauchamp et al., 2023; Chardon et al., 2023; M. Gorassini et al., 2002; Johnson et al., 2017).

Human data examining the effects of exercise training on intrinsic motoneuron properties is mixed. Studies employing subcortical stimulation of corticospinal axons (Angius et al., 2024; Ansdell et al., 2020; Nuzzo et al., 2017) and sensory activation of motoneurons (for meta-analysis, see Siddique et al., 2020) often show no changes after training, but in a cross-sectional model of chronic resistance training, alterations in the responsiveness to cervicomedullary stimulation *across contraction intensities* have been observed (Pearcey et al., 2014). Furthermore, studies examining MU discharge properties (Del Vecchio et al., 2019) and estimates of PIC magnitude (via onset-offset hysteresis of pairs of MUs; ΔF, Orssatto et al., 2023) both show increases in discharge rate and estimates of PICs following short-term training. In a cross-sectional comparison, a previous study estimated human PIC magnitude (ΔF) in several lower limb muscles of resistance-trained, endurance-trained and untrained individuals and observed no differences between the groups (Goreau et al., 2024). However, this investigation was limited to a low level of contraction force (20% of maximal voluntary force, MVF) and therefore low-threshold MUs.

Several lines of evidence suggest that adaptations in biophysical properties of motoneurons might be limited to, or be more readily apparent in, higher threshold motoneurons. For example, rat motoneuron cell capacitance has been shown to increase following endurance training (Beaumont & Gardiner, 2003), whereas the current for inducing rhythmic firing and rheobase current have been shown to decrease following weightlifting (Krutki et al., 2017) and compensatory overload (tenotomy of synergists; Krutki et al., 2015) to a greater extent or exclusively in higher threshold motoneurons. We have also shown that the excitatory-inhibitory synaptic coupling is modulated with contraction force. Specifically, inhibition patterns appear to be tonic at low forces, but more reciprocal (push-pull) in relation to excitation at higher intensities, leading to greater linearity of the ascending discharge rate and greater estimates of PIC magnitude (Škarabot et al., 2025). Given the reported changes in motoneuron receptors linked to the modulation of tonic inhibition after exercise in rodents (Woodrow et al., 2013) and the potent effect of exercise on synaptic inhibition in humans (e.g. reciprocal inhibition; Geertsen et al., 2008), the differences in intrinsic motoneuron excitability and/or motor commands are more likely to differ between trained compared to untrained individuals at higher contraction forces.

Here, we estimated potential adaptations in the intrinsic properties of human motoneurons and aimed to garner insights into the proportion of excitatory, inhibitory, and neuromodulatory motor commands by examining the MU discharge patterns during contractions up to 70% of MVF in resistance-trained, endurance-trained, and untrained individuals. We hypothesised that MU discharge patterns would become more linear and exhibit greater onset-offset discharge rate hysteresis in trained individuals at greater contraction intensities due to enhanced push-pull pattern of excitation-inhibition coupling. Additionally, we quantified MU discharge properties during a triangular contraction superimposed on a steady low-force task, which has been shown to degrade force control likely due to impaired inhibitory control of PICs (Beauchamp et al., 2025). We hypothesised that trained individuals would exhibit smaller degradation of force control due to enhanced inhibitory control.

## MATERIALS AND METHODS

### Participants

A total of 69 participants took part in this cross-sectional study (Table 1). Participants were classified as resistance-trained (n = 23, 6 females) if they reported engagement in consistent (>10 months a year), systematic, progressive resistance training for ≥ 3 years (mean ± standard deviation [SD]: 9 ± 3 years), at least 3 days a week, with at least one training session a week devoted to lower limb training. Individuals were considered endurance-trained (n = 23, 6 females) if they reported consistent engagement in systematic, progressive endurance training (running or cycling) for ≥ 3 years (mean ± SD: 10 ± 4 years), for at least 3 days a week. The remaining participants were recreationally active but did not report partaking in a systematic, progressive exercise programme and were thus classified as untrained controls (n = 23, 6 females). The physical activity levels of participants were assessed via the International Physical Activity Questionnaire (Craig et al., 2003; Table 1). The general exclusion criteria for participation in the study involved cardiovascular, neuromuscular, or musculoskeletal impairments, and taking neuroactive medication. Experimental procedures were approved by the Loughborough University ethics committee (2022-8022-12278) and were performed in accordance with the latest version of the *Declaration of Helsinki*, except for registration in database. Before participation, participants provided written, informed consent.

**Table 1.** Participant characteristics, maximal strength, and motor unit identification.

|  | Resistance-trained |  | Untrained |  | Endurance-trained |  |
| --- | --- | --- | --- | --- | --- | --- |
|  | Participant characteristics |  |  |  |  |  |
| Age (years) | 23 ± 4 |  | 24 ± 2 |  | 24 ± 6 |  |
| Mass (kg) | 83.8 ± 17.5 |  | 69.5 ± 12.1<sup>##</sup> |  | 68.1 ± 8.6<sup>###</sup> |  |
| Height (m) | 1.75 ± 0.08 |  | 1.73 ± 0.09 |  | 1.78 ± 0.09 |  |
| PA levels (MET.min/wk) | 6401 [5380, 7423]<sup>**</sup> |  | 3819 [2695, 4942] |  | 6685 [5641, 7729]<sup>**</sup> |  |
|  | Isometric strength |  |  |  |  |  |
| MVF (N) | 394 [353, 436] |  | 291 [249, 333]<sup>##</sup> |  | 337 [296, 379] |  |
|  | Motor unit identification |  |  |  |  |  |
|  | Per trial | Total | Per trial | Total | Per Trial | Total |
| Triangle 30% MVF | 20 ± 11<br>(5 – 43) | 915 | 15 ± 9<br>(3 – 42) | 700 | 25 ± 9<br>(7 – 42) | 1147 |
| Triangle 50% MVF | 18 ± 9<br>(3 – 33) | 838 | 13 ± 7<br>(2 – 30) | 598 | 21 ± 8<br>(3 – 36) | 976 |
| Triangle 70% MVF | 14 ± 7<br>(2 – 27) | 654 | 12 ± 7<br>(3 – 29) | 530 | 19 ± 8<br>(3 – 35) | 836 |
| Sombrero 10/30% MVF | 18 ± 10<br>(2 – 35) | 397 | 15 ± 10<br>(3 – 36) | 353 | 21 ± 10<br>(4 – 43) | 443 |
| Sombrero 10/50% MVF | 14 ± 8<br>(2 – 29) | 312 | 10 ± 7<br>(1 – 27) | 232 | 19 ± 8<br>(5 – 34) | 399 |
| Sustained 10% MVF | 23 ± 11<br>(6 – 45) | 533 | 19 ± 9<br>(8 – 40) | 412 | 26 ± 12<br>(4 – 50) | 554 |

Data are presented as mean ± standard deviation or estimated marginal means [95% confidence interval]. MVF = maximal voluntary force, PA = physical activity. <sup>###</sup>*p* < 0.001, <sup>##</sup>*p* < 0.01 compared to resistance-trained; <sup>\*\*</sup>*p* < 0.01 compared to untrained.

### Experimental design and protocol

Participants visited the laboratory for a familiarisation session to practice maximal and submaximal unilateral isometric contractions with ankle dorsiflexors. Participants returned to the laboratory two to ten days later to complete the experimental session. A warmup was performed initially with participants isometrically contracting their dorsiflexors for 3-5 seconds at 50 (×2), 70 (×2) and 90% (×1) of perceived maximal effort. After that, two isometric dorsiflexion contractions were performed with maximal effort (30-60 s rest between contractions). An additional trial was performed if the instantaneous peak force during the performance of maximal effort contractions differed by more than 5%. The greatest instantaneous force across the trials with maximal effort was taken as maximal voluntary force (MVF).

Participants were then tasked to perform triangular contractions with linear increases/decreases of force over 10 seconds up to/from the target of 30, 50, and 70% MVF. Two trials at each contraction level were performed, separated by 30-60 seconds of rest. Following triangular contractions, participants performed a task of a sustained contraction at 10% MVF with a superimposed triangle up to 30 and 50% MVF (Beauchamp et al., 2025; Goodlich et al., 2024; >60 seconds of rest between trials). We colloquially refer to this task as ‘sombrero’. Specifically, the force was increased over 3.3 seconds from rest to 10% MVF which was maintained for 10 seconds as accurately as possible (plateau 1; Figure 1); the force was then linearly increased/decreased over 6.7 seconds to/from the target (30 or 50% MVF), followed by another sustained phase of 10 seconds (plateau 2), and then a slow relaxation over 3.3 seconds (total contraction duration of 40 seconds). Two trials were performed at each force level; the one with the force profile that most closely matched the target based on visual inspection was selected for analysis. Finally, participants performed a sustained task of the equivalent duration to the sombrero task at 10% MVF (3.3-second increase/decrease to/from target, 33.4-second plateau).

**Figure 1.**
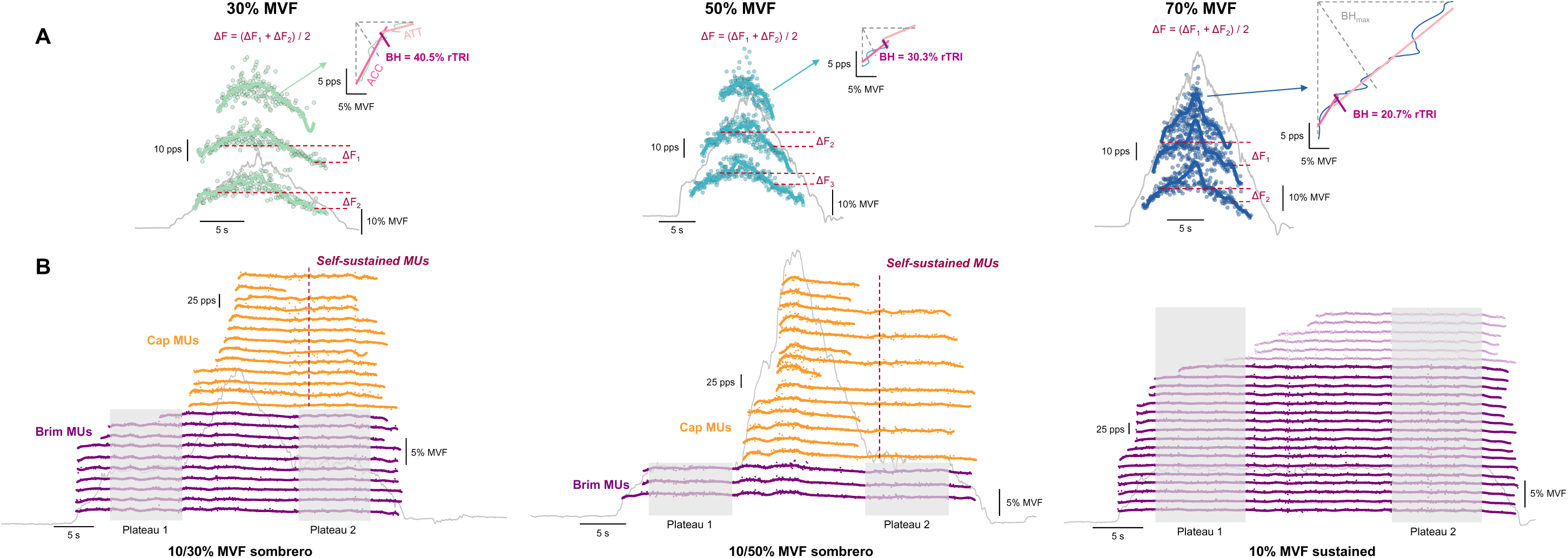
Analysis of motor unit discharge patterns and prolongation of motor unit discharge rate. A: Motor unit (MU) discharge rates during triangular contractions to 30, 50 and 70% of maximal voluntary force (MVF) were smoothed with support vector regression. We then calculated onset-offset discharge rate hysteresis (ΔF) of suitable MU pairs (only 3 units shown for clarity); the test (higher-threshold) unit ΔF values with multiple reporter (lower-threshold) units were averaged. The ascending discharge rate was expressed as a function of force and the length of the maximal orthogonal vector (brace height, BH) between the linear line between onset and peak discharge rate and the smoothed discharge rate was calculated to quantify non-linearity (normalised to the height of the right triangle, BH_max_). The point of brace height acted as a separator between the acceleration (ACC) and post-acceleration attenuation (ATT) phases for which slopes were calculated. B: the identified MUs during “sombrero” (10/30% and 10/50% MVF) and sustained (10% MVF) contractions. The units recruited during the first plateau were classified as “brim” MUs, whereas those recruited during the triangle were classified as “cap” MUs. The behaviour of brim units was compared in the first relative to the second plateau; similar analyses were performed in the sustained contraction at 10% MVF (the units had to display at least 40 discharges during the first plateau to be included; examples of excluded units are presented in opaque colour during sustained contraction). For cap MUs, we quantified the duration of MU discharge with respect to its theoretically predicted derecruitment. Further, the MUs that were discharging for longer than 2 seconds past their theoretical derecruitment were classified as self-sustained discharging MUs and were expressed as a proportion of all cap MUs.

### Experimental procedures

#### Force recordings

Ankle dorsiflexion forces were measured via a strain gauge (CCT Transducer s.a.s., Torino, Italy) attached to the ankle ergometer (NEG1, OT Bioelettronica, Torino, Italy) that was fitted to a rigid, custom-made chair upon which participants were seated during the experiment. The ankle, knee and hip were positioned at 10° of plantar flexion (0° = anatomical position), and 180° and 120° (180° = full extension), respectively. The foot of the dominant leg was strapped to the ergometer footplate with Velcro at the tarsometatarsal and metatarsal-phalangeal joints, and the knee was strapped to minimise any knee flexion. The analogue signal acquired by a strain gauge was amplified (×200, Forza-B, OT Bioelettronica) and digitally converted and sampled via an auxiliary channel of a 16-bit amplifier (Quattrocento, OT Bioelettronica).

#### High-density electromyography

Multichannel electromyogram signals were recorded from the tibialis anterior muscle with a 64-channel array electrode (13×5 electrode arrangement, 1 mm electrode diameter, 8 mm electrode distance; OT Bioelettronica). After skin preparation (shaving, abrasion and cleaning), a disposable bi-adhesive foam layer (Spes Medica, Battipaglia, Italy) with its cavities filled with conductive paste (Ten20, Weaver and Company, Aurora, CO, USA) was used to secure the placement of the electrode on the skin overlaying the TA muscle belly. A reference electrode (50×48 mm; Ambu Ltd., Cambridgeshire, UK) was placed on the medial malleolus of the ipsilateral leg, and a dampened strap electrode over the ankle of the contralateral leg was used to ground the signal. The signals were recorded in monopolar mode, band-pass filtered (10-500 Hz) and digitised using a 16-bit amplifier (Quattrocento, OT Bioelettronica).

### Data analysis

All analyses were performed offline in MATLAB (2022b, MathWorks Inc., Natick, MA USA).

#### Force signal

The voltage signal from the strain gauge was converted to force, gravity corrected, and low pass filtered (20 Hz, Butterworth, 4^th^ order) to remove the non-physiological properties.

#### High-density electromyography decomposition

Monopolar multichannel EMG signals were band-pass filtered (20-500 Hz, Butterworth, 4^th^ order). Channels of poorer quality based on area under the power spectrum were removed (>95% channels were retained), before being decomposed into individual MU spike trains with a Convolution Kernel Compensation algorithm (Holobar & Zazula, 2007). The signals from each contraction were decomposed independently, and the separation vectors initialised during the decomposition process were iteratively optimised by an experienced investigator using procedures described previously (Del Vecchio et al., 2020). During the processing of all contractions, MU spike trains were only retained if they displayed regular discharge patterns and had a pulse-to-noise ratio (Holobar et al., 2010) greater than 28 dB.

#### Motor unit discharge patterns

Motor unit discharge patterns during triangular contractions were estimated by smoothing the instantaneous discharge rates (reciprocal of the interspike interval) with support vector regression using procedures described previously (Beauchamp et al., 2022). The peak discharge rate was taken as the maximal value of the smoothed discharge rate profiles, whereas the recruitment threshold was estimated as the force (in % MVF) corresponding to the first spike in the binary spike train. We also calculated the difference between peak and initial discharge rate to estimate the magnitude of the ascending discharge rate modulation. The magnitude of PICs was estimated by quantifying the onset-offset hysteresis (ΔF; Gorassini et al., 1998) of suitable pairs of high-threshold (test) units with respect to the lower-threshold (reporter) units (Hassan et al., 2020). For test units with multiple suitable reporter units, ΔF values were averaged per test unit (A. S. Hassan et al., 2021). A suitable pair of MUs was defined according to the following criteria: 1) rate-rate correlation of r^2^ > 0.7 to increase the likelihood that the pairs of MUs received common synaptic input (Gorassini et al., 2004); 2) test unit had to be recruited at least 1 s after the recruitment of the reporter unit to increase the likelihood that PICs were fully activated (Hassan et al., 2020); and 3) the reported unit discharge rate modulation had to be greater than 0.5 pps whilst the test unit was active, to minimise the effect of saturation on ΔF estimates (Stephenson & Maluf, 2011). Additionally, to account for potential confounds when comparing differences in discharge rate hysteresis across contraction levels, we normalised ΔF estimates to the maximal discharge rate modulation of the reporter units from test-recruitment to reporter-unit derecruitment for each test-reporter unit pair (Škarabot et al., 2025; Figure 1).

The ascending MU discharge rate modulation was examined with a quasi-geometric approach. The extent of non-linearity (“brace height”), which is influenced by neuromodulation but is minimally influenced by the pattern of inhibition in computer simulations of biologically-realistic motoneuron pools (Beauchamp et al., 2023; Chardon et al., 2023; Škarabot et al., 2025), was computed as the maximal orthogonal vector between a theoretical linear line between onset and peak discharge and the ascending smoothed MU discharge rate with respect to the change in force. This vector was then normalised to the height of the right triangle with a hypothenuse between the onset and peak ascending discharge rate that represents the maximal theoretical activation of PICs (i.e. maximal non-linearity with vertical acceleration and complete post-acceleration saturation). Such normalisation accounts for the differences in relative discharge rate modulation (e.g. associated with different contraction forces). The estimation of the ascending non-linearity was considered valid in cases where the acceleration slope was positive, the normalised values did not exceed 200%, and the peak MU discharge rate occurred before peak force (Beauchamp et al., 2023). The instance of brace height on the ascending MU discharge rate modulation acted as the separator of the acceleration (secondary discharge range) and post-acceleration attenuation (tertiary discharge range) phases, for which we calculated slopes. The attenuation slope has been shown to be particularly influenced by the pattern of inhibitory synaptic input (Beauchamp et al., 2023); notably, however, this sensitivity to inhibitory gain is likely only relevant when assessed at the same contraction level due to the greater relative increase in force with respect to discharge rate when comparisons are made across contraction levels (Škarabot et al., 2025).

#### Motor unit discharge rate prolongation

To facilitate the assessment of differences between groups in MU prolongation, MUs identified during the sombrero tasks were separated into two cohorts: “brim” units, which included units recruited in the beginning, i.e. before the superimposed triangular force task; and “cap” units, which included units recruited during the superimposed triangular ramp (Figure 1B). For brim units, we quantified the mean discharge rate during the two plateaus. These calculations were additionally constrained to MUs that exhibited at least 40 discharges during the first plateau. Similar analyses were performed on MUs identified during the sustained contractions; these served as a control condition to ensure that any differences between groups in the discharge behaviour during the first vs. the second plateau were not solely a function of time-dependent adaptations in motoneuron discharge rate (e.g. spike frequency adaptation(Powers et al., 1999; Sawczuk et al., 1995). For cap units, we quantified the duration of the sustained discharge as the difference in time between derecruitment and the theoretically expected point of derecruitment (i.e. if a unit was recruited 1 second after the beginning of the superimposed triangular ramp, then the theoretically expected point of derecruitment was 1 s before the end of the ramp). Additionally, we quantified the proportion of self-sustained cap MUs with respect to the whole pool of identified cap MUs; the self-sustained cap MUs were classified as those that sustained their discharge for >2 s after the theoretically expected point of derecruitment (Beauchamp et al., 2025).

### Statistical analysis

To investigate differences between groups in participant characteristics (age, height, mass, physical activity levels), maximal unilateral isometric dorsiflexion strength, and the number of the identified MUs, a linear model was constructed with group as a factor. To assess between-group differences in MU discharge patterns across contraction levels, linear mixed-effect models were constructed with group and contraction level and their interaction as fixed effects, MU recruitment threshold as a covariate, and participant ID and contraction trial nested within participant ID as random effects (i.e. *outcome ∼ 1 + group*contraction level + MU recruitment threshold + [1 | participant ID] + [1 | contraction trial: participant ID]*). During the sombrero task, we compared the discharge rate of the “brim” units as a function of group, plateau ID and their interactions (fixed factors), with recruitment threshold of a unit as covariate, and participant ID and MU ID nested within participant ID as random intercepts (i.e. *outcome ∼ 1 + group*plateau ID + MU recruitment threshold + [1 | participant ID] + [1 | MU ID: participant ID]*). Separate models were constructed for each sombrero condition (10/30 and 10/50% MVF). Similar assessments were performed on the sustained task. Furthermore, we assessed the differences in force variability (coefficient of variation of force) on these contractions (i.e. coefficient of variation of force *∼ 1 + group*plateau ID + [1 | participant ID]*). To assess differences between groups in self-sustained discharge rate of cap units, we constructed a model with self-sustained discharge duration and proportion of sustained MUs (relative to all cap units) as outcome variables, with group as fixed effect, recruitment threshold as covariate, and participant ID as random intercept (i.e. *outcome ∼ 1 + group + MU recruitment threshold + [1 | participant ID]*).

All statistical analyses were performed in R (R studio, v 2.2, R Foundation for Statistical Computing, Vienna, Austria). Linear mixed models were constructed using the *lme4* package (Bates et al., 2015). The significance of the model with and without predictor variables was performed using a Type II Wald Chi-square test with *lmerTest* package (Kuznetsova et al., 2017). The normal distribution and homoscedasticity of model residuals were inspected using quantile-quantile and box plots. If the linear assumption of the model was not satisfied, the logarithmic (peak discharge rate during triangular contractions, all variables related to cap units during sombrero contractions), square root (absolute and normalised ΔF, acceleration and attenuation slopes), or inverse (all variables related to brim units during sombrero contractions) transformations were performed. Significant main effects and interactions were further investigated with post hoc testing of pairwise and interaction contrasts of estimated marginal means (Bonferroni’s correction for multiple comparisons) using *emmeans* package (Lenth & Lenth, 2018). To appreciate the extent of pairwise differences between groups, and the differences between the plateaus during sombrero vs. sustained task we calculated the effect sizes as the absolute difference between estimated marginal means divided by the residual standard deviation of the model (Cohen’s *d*) using *emmeans* package. Data are presented as estimated marginal means [95% confidence interval]. Significance was set at an alpha level of 0.05.

## RESULTS

### Participant characteristics, maximal strength and motor unit identification

The groups did not differ in age (χ^2^(2) = 0.4, p = 0.6938) and height (χ^2^(2) = 1.7, p = 0.1988), but had different body mass (χ^2^(2) = 9.8, p = 0.0002) and reported physical activity levels (via IPAQ; χ^2^(2) = 8.3, p = 0.0006). Specifically, resistance-trained individuals were heavier compared to endurance-trained (p = 0.0005) and untrained (p = 0.0016), and untrained individuals also reported lower physical activity levels compared to endurance-trained (p = 0.0012) and strength-trained (p = 0.0034), with no differences observed between the two trained groups (p = 0.9347 and p = 0.9206, respectively; Table 1).

The groups differed in maximal isometric dorsiflexion strength (χ^2^(2) = 6.2, p = 0.0035) with resistance-trained individuals producing greater dorsiflexion isometric forces compared to untrained (p = 0.0023, *d* = 1.03) but not endurance-trained (p = 0.1367, *d* = 0.57, Table 1). There were no differences in maximal isometric dorsiflexion strength between endurance-trained and untrained individuals (p = 0.2658, *d* = 0.57).

The total number of identified units was different between groups χ^2^(2) = 6.3, p = 0.0031), with the number being greater in endurance-trained compared to untrained (p = 0.0020, *d* = 0.93), but no significant differences were noted between resistance-trained and untrained (p = 0.1708, *d* = 0.50) or endurance-trained (p = 0.1997, *d* = 0.43; Table 1).

### Motor unit discharge rate hysteresis and the ascending discharge rate non-linearity

To determine whether the organisation of motor commands and its modulation across contraction forces differed between the groups, we performed analyses of MU discharge patterns during triangular contractions with a linearly increasing and decreasing contraction force up to/from 30, 50 and 70% MVF. The pattern of MU discharge for each of the groups at each contraction level can be appreciated by ensemble averages of MU discharge patterns depicted in Figure 2.

**Figure 2.**
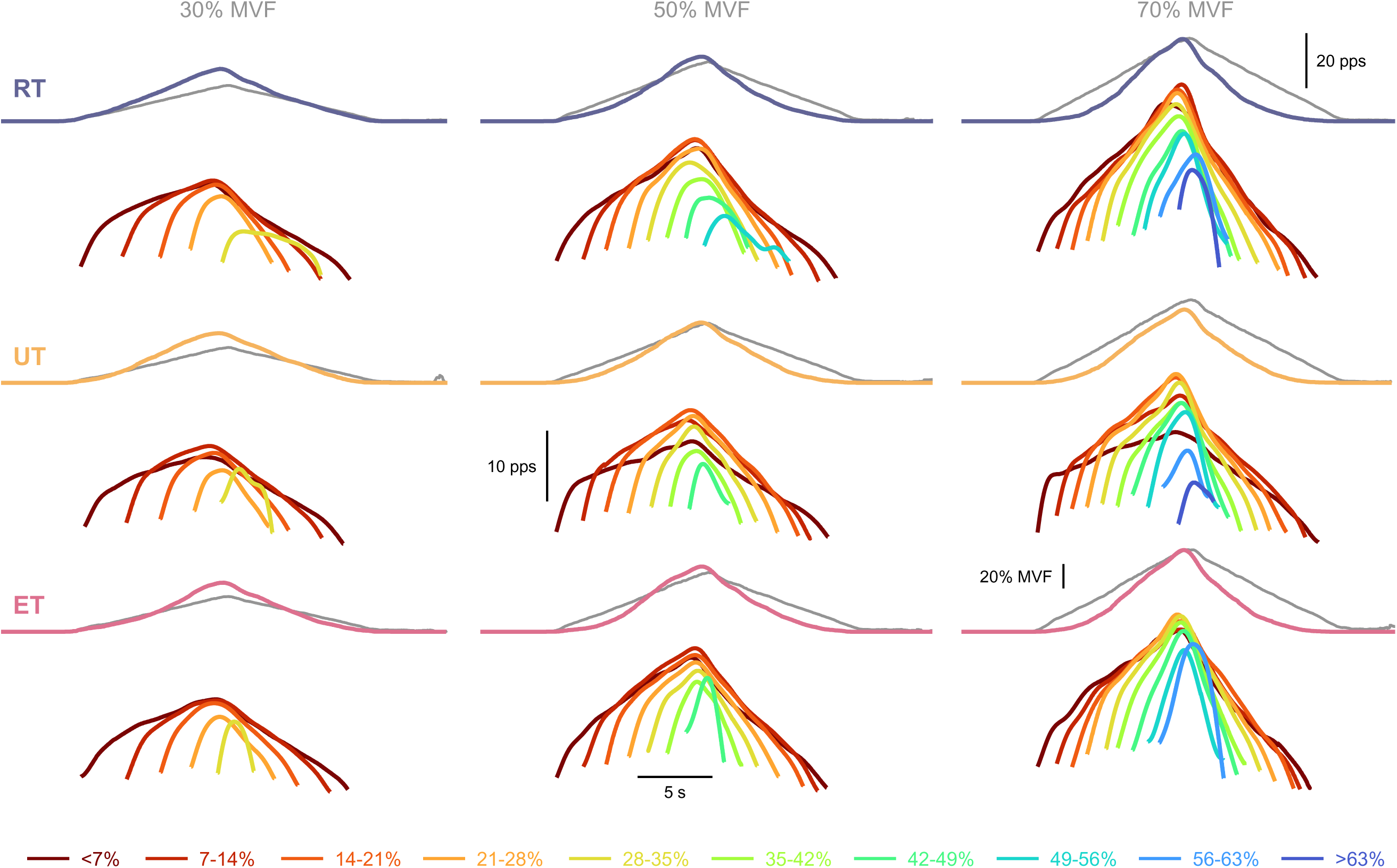
Ensemble average of motor unit discharge patterns during triangular contractions. Ensemble force traces (grey) along with cumulative smoothed discharges are depicted for each group (resistance-trained, RT; untrained, UT; endurance trained, ET) as well as ensemble averages of motor unit (MU) discharge patterns based on recruitment threshold in 7% maximal voluntary force (MVF) bins (colour-coded) during triangular contractions up to 30, 50 and 70% of MVF.

Peak discharge rate during triangular contractions was dependent on group (χ^2^(2) = 7.7, p = 0.0210), contraction level (χ^2^(2) = 9042.2, p < 0.0001) and their interaction (χ^2^(1) = 125.5, p < 0.0001). Whilst for all groups an increase in discharge rate was noted as a function of contraction force (p < 0.0001 for all), this increase was greater in resistance-trained compared to untrained from 30 to 70% (p < 0.0001) and 50 to 70% MVF (p < 0.0001), and compared to endurance-trained from 30 to 50% (p < 0.0001) and 50 to 70% MVF (p = 0.0051). Notably, the pairwise differences between groups were evident only at 50 and 70% MVF whereby resistance-(p = 0.0366, *d* = 0.63; and p = 0.0001, *d* = 1.05, respectively) and endurance-trained individuals (p = 0.0064, d = 0.78; and p = 0.0447, d = 0.60, respectively; Figure 3A) exhibited greater discharge rate compared to untrained.

**Figure 3.**
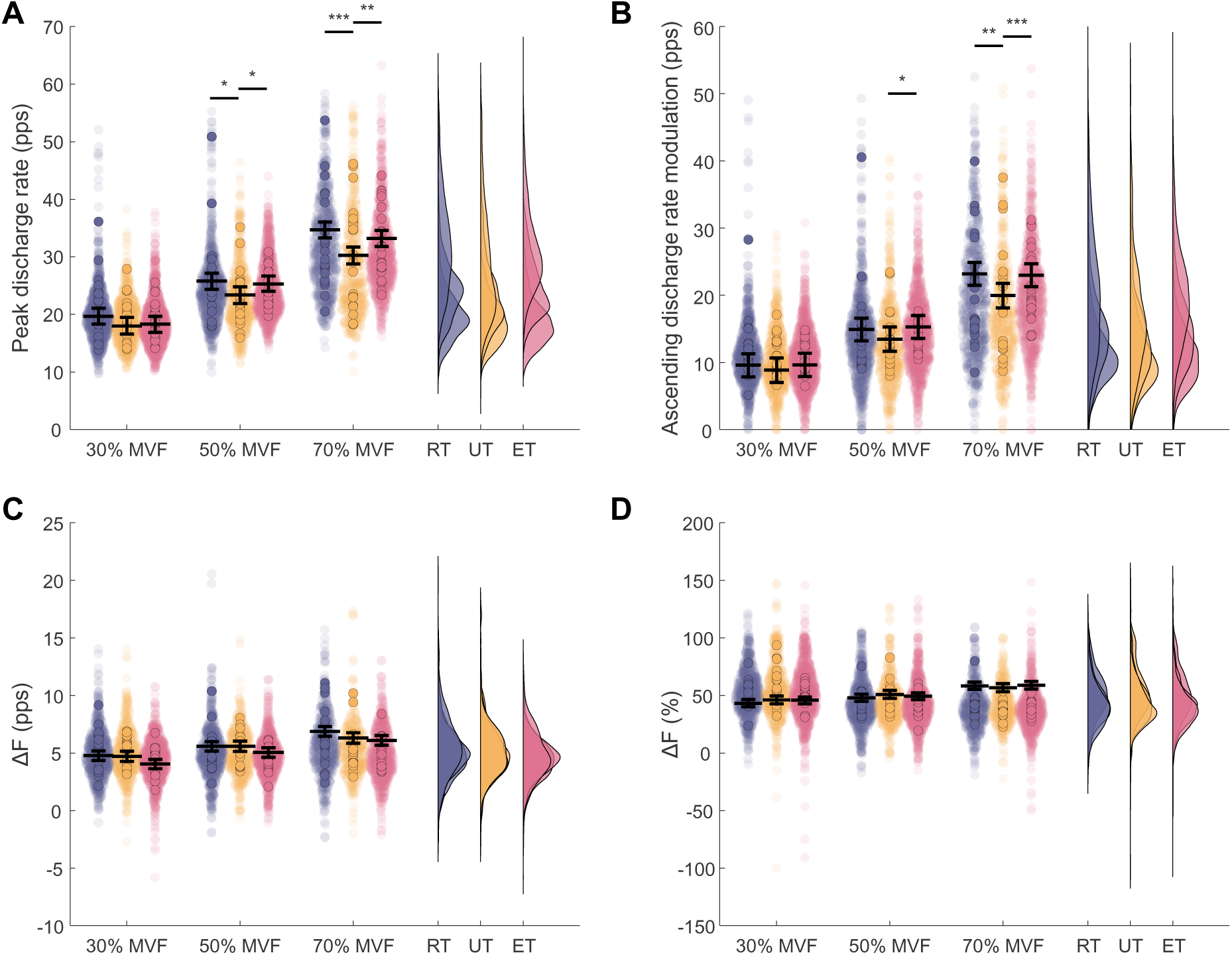
Differences in motor unit discharge rate and hysteresis between resistance-trained (RT), untrained (UT), and endurance-trained (ET) individuals during triangular contractions up to 30, 50, and 70% of maximal voluntary force (MVF) A: Peak discharge rate; B: Ascending discharge rate modulation (the difference between peak and initial discharge rate); C: motor unit (MU) discharge rate hysteresis (ΔF; in pulses per second, pps); D: MU discharge rate hysteresis normalised to the maximal discharge rate modulation for each test-reported unit pair. Individual participant and individual MU values are depicted in coloured and opaque circles, respectively, whereas the horizontal black lines denote the estimated marginal means (with 95% confidence intervals) obtained from the linear mixed statistical modelling. ***p < 0.001, **p < 0.01, *p < 0.05 pairwise differences between groups at each contraction level.

Similarly, the ascending discharge rate modulation (i.e., the difference in peak and initial discharge rate) during triangular contractions was dependent on group (χ^2^(2) = 9.0, p = 0.0112), contraction level (χ^2^(2) = 3806.2, p < 0.0001) and their interaction (χ^2^(1) = 32.8, p < 0.0001). Within each group, the ascending discharge rate modulation increased progressively with contraction force (p < 0.0001 for all), but this increase was greater in resistance-trained compared to untrained individuals between 30 and 70% (p = 0.0001) and 50 and 70% MVF (p = 0.0161). When considering pairwise comparisons, the ascending discharge rate modulation was greater in resistance-(p = 0.0024, *d* = 0.62) and endurance-trained (p = 0.0002, *d* = 0.52) at 70% MVF, and endurance compared to untrained at 50% MVF (p = 0.0056, *d* = 0.47; Figure 3B).

Discharge rate hysteresis (i.e., ΔF) was influenced by contraction level (χ^2^(2) = 523.4, p < 0.0001) and the interaction between group and contraction level (χ^2^(1) = 14.0, p = 0.0073). In all groups, discharge rate hysteresis increased with contraction force (p < 0.0001 for all), but the resistance-trained individuals exhibited a greater gain (i.e., a relative increase) in ΔF from 30 to 70% (p = 0.0091) and 50 to 70% MVF (p = 0.0169) compared to untrained individuals. However, pairwise between-group differences were not observed at any contraction level (p ≥ 0.0799, *d* ≤ 0.37; Figure 3C).

When expressed relative to the maximal theoretical hysteresis of the test unit, ΔF was predicted by contraction level (χ^2^(2) = 305.2, p < 0.0001) and the interaction between group and contraction level (χ^2^(1) = 9.5, p = 0.0498). In all groups, the normalised ΔF increased with contraction force (p ≤ 0.0001), but the interaction contrasts suggested this effect was similar across all groups (p ≥ 0.0729), and no pairwise between-group differences were detected at any contraction level (p ≥ 0.3539, *d* ≤ 0.01; Figure 3D).

The non-linearity in discharge rate modulation during the ascending phase of triangular contractions was dependent on group (χ^2^(2) = 19.6, p < 0.0001), contraction level (χ^2^(2) = 408.4, p < 0.0001) and their interaction (χ^2^(1) = 19.4, p = 0.0006). In all groups, the ascending discharge rate modulation became more linear with respect to force as contraction force increased (p ≤ 0.0001), however, this decrease was of greater magnitude in resistance-trained compared to untrained from 30 to 70% MVF (p = 0.0354) and from 30 to 50% MVF (p = 0.0298), and compared to endurance-trained from 30 to 50% MVF (p = 0.0040). When considering pairwise comparisons between groups, the ascending discharge rate modulation was more linear in resistance- and endurance-trained compared to untrained at 50% (p = 0.0001, *d* = 0.54; and p < 0.0001, *d* = 0.35, respectively) and 70% MVF (p < 0.0001, *d* = 0.57; and p = 0.0123, *d* = 0.56, respectively), whereas at 30% the ascending discharge rate was more linear in endurance-(p = 0.0068, *d* = 0.36), but not resistance-trained (p = 0.0529, *d* = 0.28) compared to untrained (Figure 4A).

**Figure 4.**
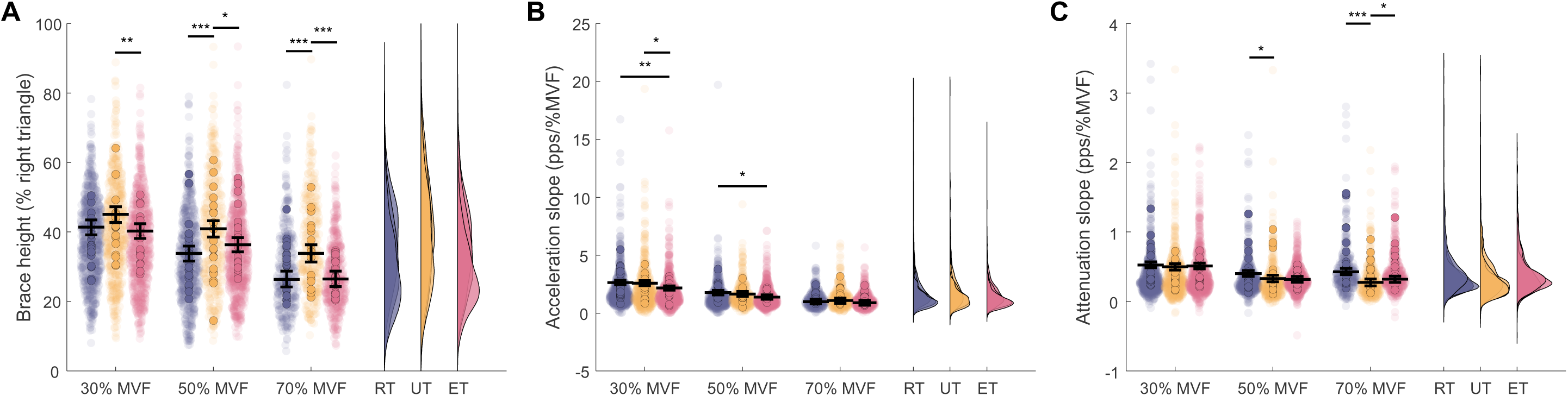
Differences in motor unit discharge patterns between resistance-trained (RT), untrained, and endurance-trained (ET) individuals during triangular contractions up to 30, 50, and 70% of maximal voluntary force (MVF) A: The ascending MU discharge rate non-linearity (brace height; as % of the right triangle), B: the slope of the MU acceleration phase; C: the slope of the post-acceleration attenuation phase. Individual participant and individual MU values are depicted in coloured and opaque circles, respectively, whereas the horizontal black lines denote the estimated marginal means (with 95% confidence intervals) obtained from the linear mixed statistical modelling. ***p < 0.001, **p < 0.01, *p < 0.05 pairwise differences between groups at each contraction level.

The discharge rate acceleration slope was predicted by group (χ^2^(2) = 8.6, p = 0.0134), contraction level (χ^2^(2) = 750.8, p < 0.0001) and their interaction (χ^2^(1) = 12.5, p = 0.0138). The acceleration slope decreased in all groups with contraction level (p < 0.0001), however, this decrease was of greater magnitude from 30 to 70% in endurance-compared to resistance-trained individuals (p = 0.0032). Endurance-trained individuals also displayed smaller acceleration slopes compared to resistance-trained and untrained individuals at 30% MVF (p = 0.0020, *d* = 0.43; and p = 0.0245, *d* = 0.34, respectively), and compared to resistance-trained at 50% MVF (p = 0.0264, *d* = 0.33; Figure 4B).

The post-acceleration discharge rate modulation (i.e., attenuation) slope was also predicted by group (χ^2^(2) = 7.8, p = 0.0202), contraction level (χ^2^(2) = 306.4, p < 0.0001) and their interaction (χ^2^(1) = 36.2, p < 0.0001). In all groups, the attenuation slope decreased from 30 to 50% (p < 0.0001) and 30 to 70% (p < 0.0001), however, it only decreased from 50 to 70% MVF in the untrained group (p = 0.0459). The interaction contrasts suggested that untrained individuals exhibited a greater relative decrease in attenuation slope compared to resistance trained during all contraction levels (p ≤ 0.0470), and similarly, the endurance-trained individuals exhibited a greater relative decrease in attenuation slope compared to resistance-trained from 30 to 50% (p = 0.0016) and 30 to 70% MVF (p = 0.0006). Consequently, the pairwise between-group differences were evident at 70% MVF where resistance-trained individuals displayed greater attenuation slopes compared to untrained (p < 0.0001, *d* = 0.67) and endurance-trained (p = 0.0332, *d* = 0.34); the endurance-trained individuals also demonstrated greater attenuation slopes compared to untrained (p = 0.0477, *d* = 0.33). Furthermore, at 50% MVF, the attenuation slopes of resistance-trained individuals were also greater compared to untrained (p = 0.0127, *d* = 0.38; Figure 3F).

Collectively, the results obtained from MU discharge rates during triangular contractions suggest that the trained groups had greater peak discharge rates and greater ascending discharge rate modulation compared to untrained, but these differences were more evident at greater contraction levels. The greater gain of peak discharge rates and ascending discharge rate modulation across contraction intensities in the trained groups was accompanied by differences in the linearity of the ascending MU discharge rate. In trained individuals, ascending discharge rate modulation was more linear, with greater attenuation slopes compared to untrained. Additionally, compared to untrained, resistance-trained individuals exhibited greater acceleration slopes during lower contraction forces (Table 2).

**Table 2.** Summary of the comparison of discharge rate characteristics between resistance-, endurance-, and untrained individuals during triangular contractions at 30, 50 and 70% of maximal voluntary force.

|  | RT |  |  | UT |  |  | ET |  |  |
| --- | --- | --- | --- | --- | --- | --- | --- | --- | --- |
|  | 30% | 50% | 70% | 30% | 50% | 70% | 30% | 50% | 70% |
| Peak discharge rate (pps) | ↔ | >UT | >UT | ↔ | <RT, <ET | <RT, <ET | ↔ | >UT | >UT |
| Ascending discharge rate modulation (pps) | ↔ | ↔ | >UT | ↔ | <ET | <RT, <ET | ↔ | >UT | >UT |
| $\Delta F$ (pps) | ↔ | ↔ | ↔ | ↔ | ↔ | ↔ | ↔ | ↔ | ↔ |
| $\Delta F$ (%) | ↔ | ↔ | ↔ | ↔ | ↔ | ↔ | ↔ | ↔ | ↔ |
| Brace height (%rTRI) | ↔ | >UT | >UT | <ET | <RT, <ET | <RT, <ET | >UT | >UT | >UT |
| Acceleration slope (pps/%MVF) | >UT, >ET | >ET | ↔ | <RT | ↔ | ↔ | <RT | <RT | ↔ |
| Attenuation slope (pps/%MVF) | ↔ | >UT | >UT | ↔ | <RT | <RT, <ET | ↔ | ↔ | >UT |

*RT = resistance-trained, UT = untrained, ET = endurance-trained; MVF = maximal voluntary force; pps = pulses per second; rTRI = right triangle.*

### Motor unit discharge rate prolongation during sombrero contractions

#### Cap Units

The duration of MU discharge rate prolongation was not different between groups either for the 10/30% (χ^2^(2) = 2.9, p = 0.2311) or 10/50% MVF sombrero contractions (χ^2^(2) = 1.6, p = 0.4413; Figure 5A). Similarly, the groups did not differ in the proportion of sustained units during both the 10/30 (χ^2^(2) = 4.3, p = 0.5024) and 10/50% MVF sombreros (χ^2^(2) = 1.7, p = 0.4175; Figure 5B).

**Figure 5.**
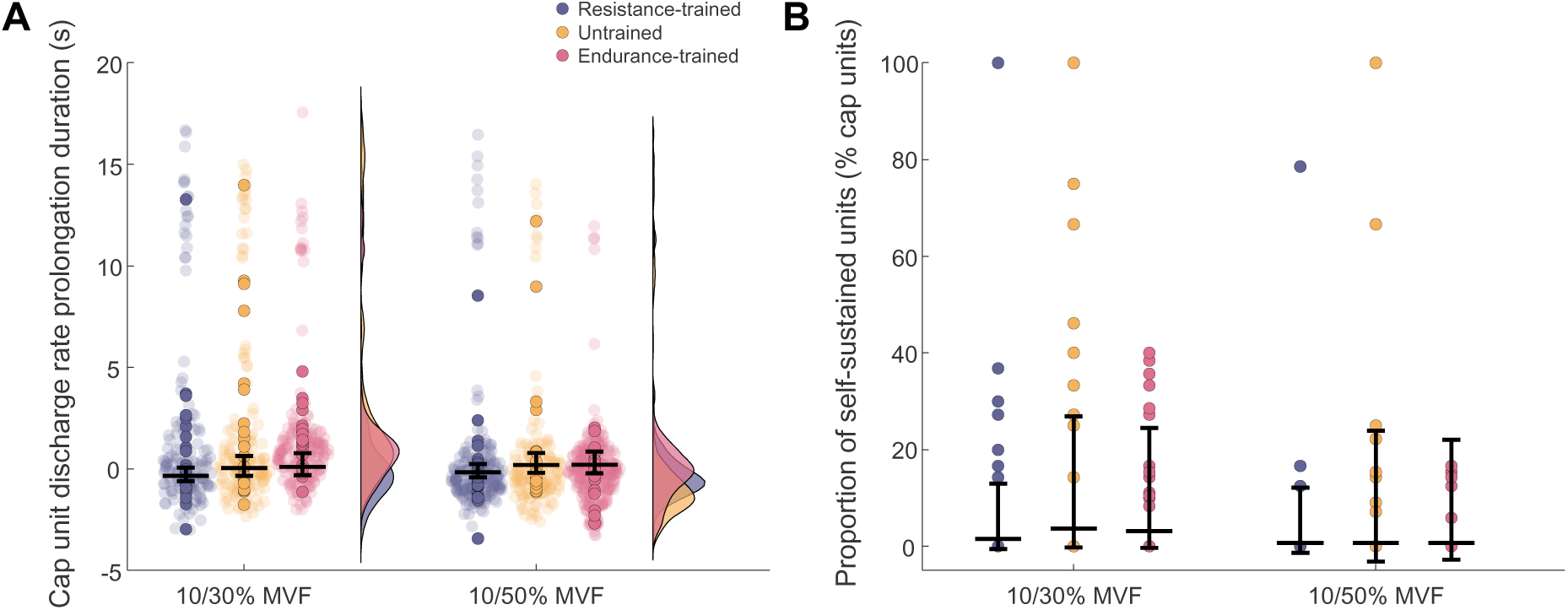
Motor unit discharge characteristics of cap units during “sombrero” contractions. To facilitate analysis of motor unit (MU) discharge rate prolongation during “sombrero” contractions, the units were clustered into “brim” units (i.e. those recruited on the first plateau of the steady portion of the task), and “cap” units (i.e. those recruited during the triangular ramp superimposed onto sustained contraction. A: Duration of discharge rate prolongation of cap units with respect to the theoretical derecruitment of the respective unit. B: Proportion of self-sustained units relative to the entire identified pool of cap units (n.b. a unit was considered to exhibit self-sustained discharge rate if it was active >2 seconds after its theoretical derecruitment). Individual participant and individual MU values are depicted in coloured and opaque circles, respectively, whereas the horizontal black lines denote the estimated marginal means (with 95% confidence intervals) obtained from the linear mixed statistical modelling.

#### Brim units

The discharge rate during 10/30 and 10/50% sombrero contractions differed between plateaus (10/30% MVF: χ^2^(1) = 1508.0, p < 0.0001, *d* = 2.37, 10/50% MVF: χ^2^(1) = 682.2, p < 0.0001, *d* = 2.77), but not groups (χ^2^(2) = 1.1, p = 0.5815). However, there was a significant interaction between group and plateau (10/30% MVF: χ^2^(2) = 15.7, p = 0.0004; 10/50% MVF: χ^2^(2) = 10.8, p = 0.0046). Specifically, MU discharge rate decreased between the two plateaus in all groups for both 10/30 and 10/50% MVF conditions (p < 0.0001 for all), however, this decrease was greater in untrained compared to endurance-trained during the 10/30% (p = 0.0005) and 10/50% MVF sombrero (p = 0.0094), and in resistance-compared to endurance-trained at 10/30% MVF (p = 0.0174; Figure 6A).

**Figure 6.**
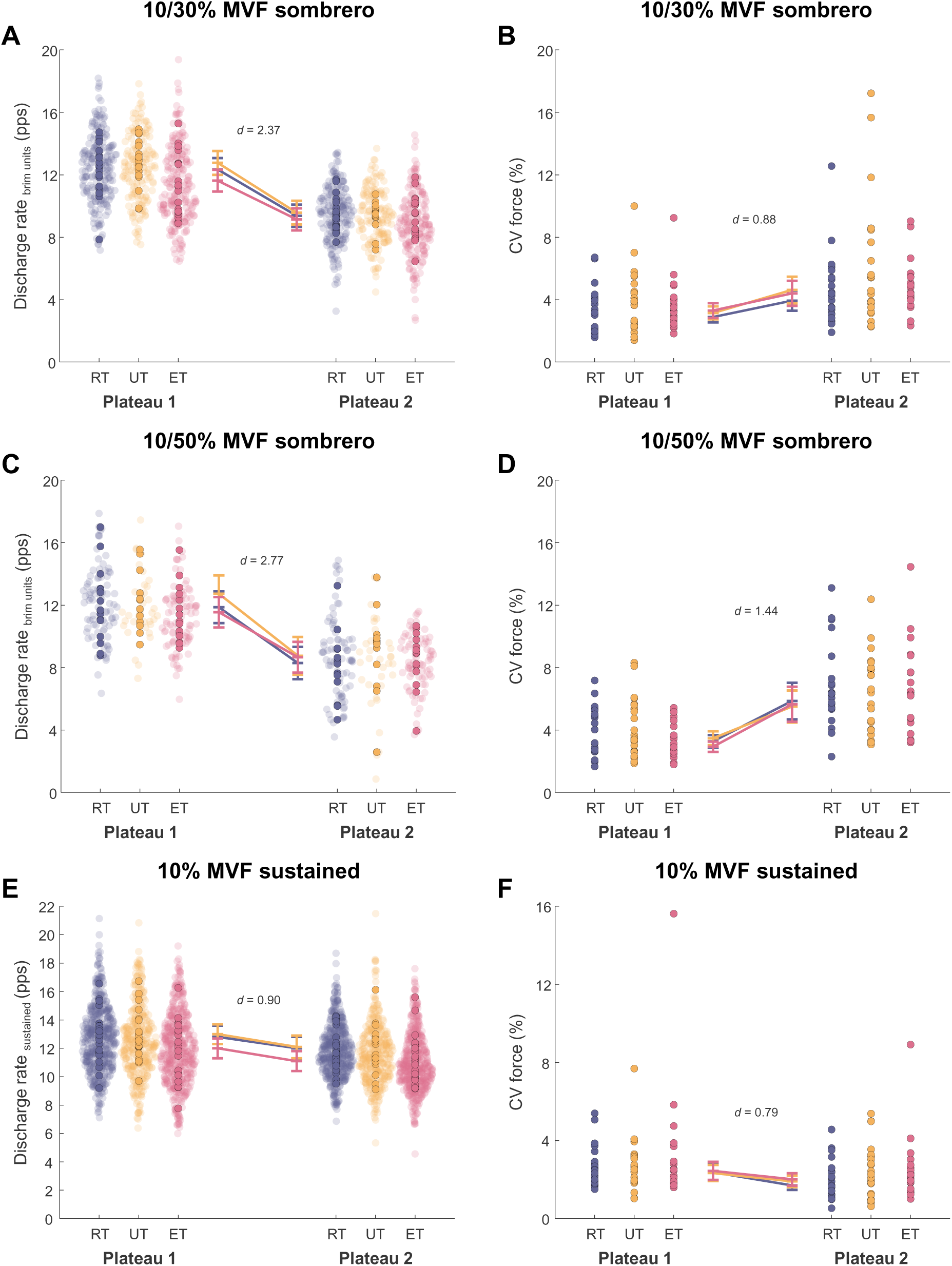
Motor unit discharge characteristics of “brim” units during “sombrero” and sustained contractions. To facilitate analysis of motor unit (MU) discharge rate characteristics during “sombrero” contractions, the units were clustered into “brim” units (i.e. those recruited on the first plateau of the steady portion of the task), and “cap” units (i.e. those recruited during the triangular ramp superimposed onto sustained contraction). During sustained contractions, MU discharge characteristics were quantified in the time periods equivalent to the plateaus during sombrero contractions. A, C: Discharge rate of brim units during the first and second plateau of the steady part of the task during 10/30% and 10/50% of maximal voluntary force (MVF) sombreros, respectively. B, D: Coefficient of variation of force during the first and second plateau of the steady part of the task during 10/30% and 10/50% MVF sombreros, respectively. E: Discharge rate of units during sustained contractions at 10% MVF in the time period equivalent to the plateaus during sombreros. F: Coefficient of variation of force during sustained contractions at 10% MVF in the time period equivalent to the plateaus during sombreros. Individual participant and individual MU values are depicted in coloured and opaque circles, respectively, whereas the horizontal black lines denote the estimated marginal means (with 95% confidence intervals) obtained from the linear mixed statistical modelling. The effect sizes shown (Cohen’s d; the absolute difference between estimated marginal means divided by the residual standard deviation of the model) were constructed to characterise the differences between the plateaus.

#### Control of force output

The variability in force production (coefficient of variation of force) did not differ between groups (10/30% MVF: χ^2^(2) = 1.9, p = 0.3840; 10/50% MVF: χ^2^(2) = 1.3, p = 0.5142), but was dependent on plateau (10/30% MVF: χ^2^(1) = 25.8, p < 0.0001, d = 0.88; 10/50% MVF: χ^2^(1) = 67.6, p < 0.0001, d = 1.44). In all groups, the variability of force production increased in the second compared to the first plateau during both 10/30% (p ≤ 0.0041) and 10/50% (p ≤ 0.0003) sombrero contractions (Figure 6B).

#### Sustained contractions

The discharge rate during sustained contractions in periods equivalent to the plateau during sombrero contractions was different (i.e., decreased; χ^2^(1) = 525.8, p < 0.0001, *d* = 0.90), but there was no main effect of group (χ^2^(2) = 3.6, p = 0.1631; Figure 6C). Similarly, the coefficient of variation of force was not different between groups (χ^2^(2) = 0.8, p = 0.6753). However, unlike in sombrero contractions, the force became less variable between the time periods equivalent to the plateau during sombrero contractions (χ^2^(1) = 19.7, p < 0.0001, *d* = 0.79; Figure 6D).

## DISCUSSION

In this study, we estimated the intrinsic properties of human motoneurons and garnered insights about the proportion of excitatory, inhibitory, and neuromodulatory motor commands in chronically resistance and endurance trained in comparison with untrained individuals to determine potential long-term training adaptations. In agreement with our hypothesis, in both resistance- and endurance-trained individuals, discharge patterns at high contraction forces reached greater rates, were more linear, and had steeper post-acceleration slopes compared to untrained. This either suggests a more reciprocal (push-pull) pattern of synaptic inhibition-excitation coupling or greater overall excitatory synaptic input to the motor pool that possibly contributes to greater MU discharge rates and ascending discharge rate modulation. Nevertheless, and in contrast to our hypothesis, there was no difference in task performance or the underlying metrics of intrinsic motoneuron excitability during sombrero contractions. Finally, though onset-offset hysteresis was not different between groups (i.e., suggesting similar contribution of PICs to MU discharge rate prolongation), resistance-compared to endurance-trained individuals exhibited greater acceleration slopes at lower contraction levels, suggesting greater contribution of neuromodulatory input, which enhances PICs, to MU discharge rate acceleration in lower-threshold MUs.

### Greater discharge rates in trained individuals are accompanied by more linearity during higher contraction forces

During triangular contractions, both groups of trained individuals exhibited greater MU discharge rates compared to untrained, however, this between-group difference was limited to higher contraction forces (≥50% MVF). This observation contrasts with prior human studies that have suggested no differences in MU discharge rates, even at higher contraction forces, in biceps brachii and the vastii muscles (Casolo et al., 2021; Škarabot et al., 2023). This discrepancy might be related to methodological factors such as the choice of the task; unlike in the present study that used triangular contractions, others have used trapezoidal contractions with a prolonged plateau at a target force level where alterations in motoneuron intrinsic properties (e.g. spike frequency adaptation; Powers et al., 1999; Vandenberk & Kalmar, 2014) could have influenced the calculation of the mean discharge rate. Notably however, our results of between-group differences in MU discharge rate being limited to higher contraction forces agree with the premise suggested by rodent studies where training-related alterations of biophysical properties of motoneurons appear more pronounced or exclusively apparent in higher threshold motoneurons (Beaumont & Gardiner, 2003; Krutki et al., 2015, 2017).

The greater MU discharge rate and ascending discharge rate modulation of trained compared to untrained individuals at higher contraction forces was accompanied by more linear ascending MU discharge patterns. The activation of PICs typically leads to distinct non-linearities in the motoneuron discharge patterns during linear increases in synaptic input, hallmarked by an initial acceleration, followed by post-acceleration rate attenuation and prolongation of MU discharge rate resulting in onset-offset hysteresis (Beauchamp et al., 2023; Heckman & Enoka, 2012; Khurram et al., 2022). More linear ascending MU discharge patterns of the trained groups indicates a smaller *relative* contribution of PICs to the ascending motoneuron discharge rate with greater excitatory synaptic input (Škarabot et al., 2025). Our results further suggest that the more linear MU discharge patterns in trained individuals are, at least partially, due to steeper post-acceleration attenuation slopes, suggesting a more reciprocal (push-pull) excitation-inhibition coupling (Beauchamp et al., 2023; Chardon et al., 2023). This effect appears more pronounced at higher contraction forces, where the pattern inhibition relative to excitation has previously been shown to shift from tonic to reciprocal/push-pull (Škarabot et al., 2025). Therefore, in trained individuals at higher forces, more reciprocal excitation-contraction coupling might reflect greater net excitatory synaptic input to motor pool, reflected in greater relative increase (i.e. gain) in MU discharge rate with contraction force, and ultimately greater production of absolute muscle forces. An alternative explanation for these results could be that excitatory synaptic input to the motor pool is enhanced with training. Though the specificity of these synaptic input modulations cannot be ascertained from our data directly, there is indirect evidence supporting training adaptations that could lead to reduced inhibitory (decreased antagonist coactivation or reciprocal inhibition; Balshaw et al., 2019; Geertsen et al., 2008; Pearcey et al., 2014) and/or enhanced excitatory synaptic input (e.g. reticulospinal excitability; Akalu et al., 2024; Glover & Baker, 2020; Glover & Baker, 2022) to the agonist motor pool to support greater force production.

Whilst exercise-induced alterations in dendritic PICs of interneurons have been observed in rodents (Chen & Dai, 2022; Ge & Dai, 2020), we did not observe any significant differences in the MU onset-offset hysteresis (an estimate of PIC magnitude) between the groups, even when accounting for differences in descending discharge rate modulation (Škarabot et al., 2025). A prior study found no differences in onset-offset hysteresis between resistance-, endurance- and untrained individuals during contraction to 20% MVF (Goreau et al., 2024), and our findings extend these observations during contractions up to 70% MVF. Though these findings might imply similar neuromodulatory input between groups, in apparent disagreement with rodent studies showing greater activity of 5-HT neurons in raphe nuclei in response to training (Ge & Dai, 2020; Ji et al., 2017), it is worth noting that motoneuron onset-offset hysteresis is dependent on both neuromodulatory input and the pattern of inhibition relative to excitation (Beauchamp et al., 2023; Chardon et al., 2023; Škarabot et al., 2025). Given the apparent differences in excitation-inhibition coupling (i.e. more reciprocal/push-pull in trained individuals), the lack of differences in onset-offset hysteresis points to the greater relative contribution of greater modulation of excitatory, rather than inhibitory input of trained individuals.

Our findings suggests that there are clear differences between trained and untrained individuals in MU discharge patterns. However, the differences between resistance and endurance-trained individuals appear more subtle. For example, whilst MU discharge patterns became more linear with greater contraction forces in both groups, this linearisation was of greater magnitude in resistance-compared to endurance-trained (as well as untrained), possibly due to resistance training providing a more potent stimulus for motoneuron adaptations at high forces. Moreover, resistance-compared to endurance-trained individuals also exhibited steeper acceleration (secondary range) slopes of MU discharge, suggesting larger PICs produced by low-voltage activation either *subthreshold* or near the onset of MU recruitment (Afsharipour et al., 2020; Bennett et al., 1998; Lee & Heckman, 1998). Such an effect has been suggested to be more prominent in low-threshold units (Afsharipour et al., 2020; Mohammadalinejad et al., 2024), and consistent with this, we demonstrate that unlike for other metrics, the differences between the two groups of trained individuals in acceleration slope were only evident at lower contraction forces (≤50% MVF). Alternatively, greater acceleration slopes of resistance-trained individuals could stem from slow inactivation of Kv1.2 channels in the juxtanodal domain of axons (Bos et al., 2021) that, in conjunction with PICs, can accelerate discharge rate with increased synaptic input (Mohammadalinejad et al., 2024).

### Sombrero contraction-induced degradations in force control are not different between trained and untrained individuals

Despite the apparent differences in modulation of inhibition-excitation coupling gleaned from MU discharge patterns during triangular contractions, there were no differences in MU discharge properties or force control during the sombrero contractions. Specifically, in this task, upon the return to a steady hold following a superimposed triangular contraction, many units recruited during the superimposed triangle exhibit self-sustained discharge, leading to impeded force control, likely due to impaired PIC inactivation (Beauchamp et al., 2025; Goodlich et al., 2022). We found no differences between groups in both force control and MU discharge properties during this task. The lack of differences in self-sustained discharge of cap units (i.e. those recruited during the superimposed triangular contractions) is consistent with the lack of between-group differences in onset-offset hysteresis, indicating training is less likely to induce adaptations affecting PIC-mediated discharge rate prolongation or inhibitory control of PICs during sustained tasks. The latter further provides evidence of a greater likelihood of altered excitation patterns contributing to more reciprocal excitation-inhibition coupling of trained individuals at greater forces.

### Methodological considerations

This study assessed the potential effect of training across contraction forces on motor commands to, and intrinsic properties of, motoneurons using a cross-sectional design. Though attempts were made to ensure a sample of highly trained individuals with consistent training over many years (see ‘Participant’ section of the Methods), the cross-sectional design inherently lacks control over the precise training practices of individuals involved in the study. Indeed, specificity of training is an important variable when considering physiological adaptations to training (Del Vecchio et al., 2024). Nevertheless, our data suggests that chronic training, and less the type of training, exerts profound effects on motoneuron behaviour during isometric contractions, and future studies should explore these physiological mechanisms in the context of training specificity and different tasks, including those that are currently less conducive for human MU recordings (e.g., dynamic tasks).

We assessed MU discharge patterns and discharge characteristics during triangular and sombrero contractions to various target forces, but the signals obtained during these contractions were decomposed individually, likely leading to MU replicates across contraction forces without our ability to ascertain the extent of them. However, we have previously demonstrated similar modulation of MU discharge patterns across contraction levels with and without motor unit tracking in the tibialis anterior muscle (Škarabot et al., 2025), suggesting that the absence of tracking herein was unlikely to have influenced our results. The inherent bias to blind source separation algorithms towards higher threshold units (Francic & Holobar, 2021) might be viewed favourably in this regard as it suggests a greater likelihood of a proportion of unique identified MUs at each contraction level. Conversely, this bias is perhaps a disadvantage for superimposition tasks such as sombrero contractions where cap units are more likely to be identified than those classified as brim units. Nevertheless, the number of identified MUs during sombrero contractions was consistent with prior studies that employed such tasks (Beauchamp et al., 2025; Goodlich et al., 2024). The potential decomposition biases are accounted for in some variables inherently (i.e. a proportion, rather than an absolute number of, units with self-sustained discharge) or by using recruitment threshold of the identified MUs as a covariate in the statistical model.

## Conclusion

The comparison of tibialis anterior motor unit discharge patterns during isometric contractions up to 70% of maximum between resistance-trained, endurance-trained and untrained individuals revealed that the trained groups modulated MU discharge rate to a greater extent, reaching greater peak discharge rates during higher contraction forces. These differences in MU discharge rate in trained individuals were also accompanied by more linear discharge patterns with greater post-acceleration rate attenuation slopes, suggesting smaller *relative* contribution of PICs to ascending discharge rate modulation. These findings can be explained by either more reciprocal/push-pull excitation-inhibition coupling or enhanced excitatory synaptic input which could both result in greater *net excitation* to the motor pool and greater force production of trained individuals.

## Acknowledgements

The authors thank Vedika Mohite and Yanbin Zhang for their assistance with participant recruitment and data collection. J.Š. was supported by Versus Arthritis Foundation Fellowship (reference: 22569). G.E.P.P. was supported by the Natural Sciences and Engineering Research Council of Canada (Discovery Grant RGPIN-2023-05862, and Discovery Launch Supplement DGECR-2023-00279).

## Conflict of interest

The authors declare no competing financial interests.

## Notes

### Competing Interest Statement

The authors have declared no competing interest.

